# Cross-reactive immunity drives global oscillation and opposed alternation patterns of seasonal influenza A viruses

**DOI:** 10.1101/226613

**Authors:** Lorenzo Gatti, Jitao David Zhang, Maria Anisimova, Martin Schutten, Ab Osterhaus, Erhard van der Vries

## Abstract

Several human pathogens exhibit distinct patterns of seasonality and circulate as pairs of discrete strains. For instance, the activity of the two co-circulating influenza A virus subtypes oscillates and peaks during winter seasons of the world’s temperate climate zones. These periods of increased activity are usually caused by a single dominant subtype. Alternation of dominant strains in successive influenza seasons makes epidemic forecasting a major challenge. From the start of the 2009 influenza pandemic we enrolled influenza A virus infected patients (*n* = 2,980) in a global prospective clinical study. Complete hemagglutinin (HA) sequences were obtained from 1,078 A/H1N1 and 1,033 A/H3N2 viruses and were linked to patient data. We then used phylodynamics to construct high resolution spatio-temporal phylogenetic HA trees and estimated global influenza A effective reproductive numbers (*R*) over time (2009-2013). We demonstrate that *R*, a parameter to define host immunity, oscillates around *R* = 1 with a clear opposed alternation pattern between phases of the A/H1N1 and A/H3N2 subtypes. Moreover, we find a similar alternation pattern for the number of global virus migration events between the sampled geographical locations. Both observations suggest a between-strain competition for susceptible hosts on a global level. Extrinsic factors that affect person-to-person transmission are a major driver of influenza seasonality, which forces influenza epidemics to coincide with winter seasons. The data presented here indicate that also cross-reactive host immunity is a key intrinsic driver of global influenza seasonality, which determines the outcome of competition between influenza A virus strains at the onset of each epidemic season.

**Significance statement:** Annual influenza epidemics coincide with winter seasons in many parts of the world. Environmental factors, such as air humidity variation or temperature change, are commonly believed to drive these seasonality patterns. Interestingly, three out of the four latest pandemics (1918, 1968 and 2009) did not spread in winter initially, but during summer. This questions to what extent other factors could also impact virus spread among humans. We demonstrate that cross-reactive host immunity is a key factor. It drives the well-known seasonal patterns of virus activity oscillation and alternation of the dominant influenza virus subtype in successive seasons. Furthermore, this factor may also explain the efficient spread of pandemic viruses during summer when cross-reactive host immunity is relatively low.

## Introduction

Several human respiratory viruses circulate as groups of discrete pathogenic entities exhibiting distinct patterns of seasonality (1, 2). For influenza virus such patterns have been studied extensively (3-5). In the world’s temperate climate zones influenza activity oscillates and synchronizes with winter periods, while in tropical regions activity appears to be year-around or split into different seasons (4). They have been attributed largely to ‘extrinsic’ factors driving efficient virus spread (6), like air humidity variations (7), seasonal influences on host susceptibility (8), and societal structure and behavioural patterns (9). Susceptible-Infection-Recovery (SIR) epidemiological modelling predicted that also cross-reactive immunity between subtypes plays a role (10-13). Such ‘intrinsic’ factor may also be attributed to other aspects of influenza epidemiology, like the replacement of a seasonal strain by a pandemic virus. This occurred for the last time during the 2009 influenza pandemic when the seasonal A/H1N1 was replaced by the pandemic A/H1N1 virus. Interestingly, like the 1918 and 1968 pandemics this virus did not spread in winter, but during the 2009 northern hemisphere (NH) summer.

To date, the newly introduced pandemic 2009 A/H1N1 virus continues to co-circulate with the A/H3N2 subtype causing seasonal epidemics in humans. Both influenza A viruses are under intense selective pressure by the host immune system and they continuously evolve to persist in humans. Viruses escape from pre-existing immunity through mutation at antigenic sites at the globular head of the hemagglutinin (HA). This is a major virus surface glycoprotein and primary target of host neutralizing antibodies. Continual viral presence in the population on the other hand results in a ‘landscape of immunity’ (11, 14), which new ‘antigenic drift’ viruses need to overcome to fuel new epidemics. A typical phylogenetic tree of HA is shaped, as a result of this cat-and-mouse game, into a single trunk tree with short-lived branches (15) (Fig. 1). Virus strains that are antigenically similar cluster along the trunk of the tree with only a limited number of amino acid positions involved in the jump from an existing into a new antigenic cluster (16). These positions were previously identified with data obtained from the hemagglutination inhibition (HAI) assay, a serological test to assess neutralizing antibody responses to HA.

**Fig. 1.**
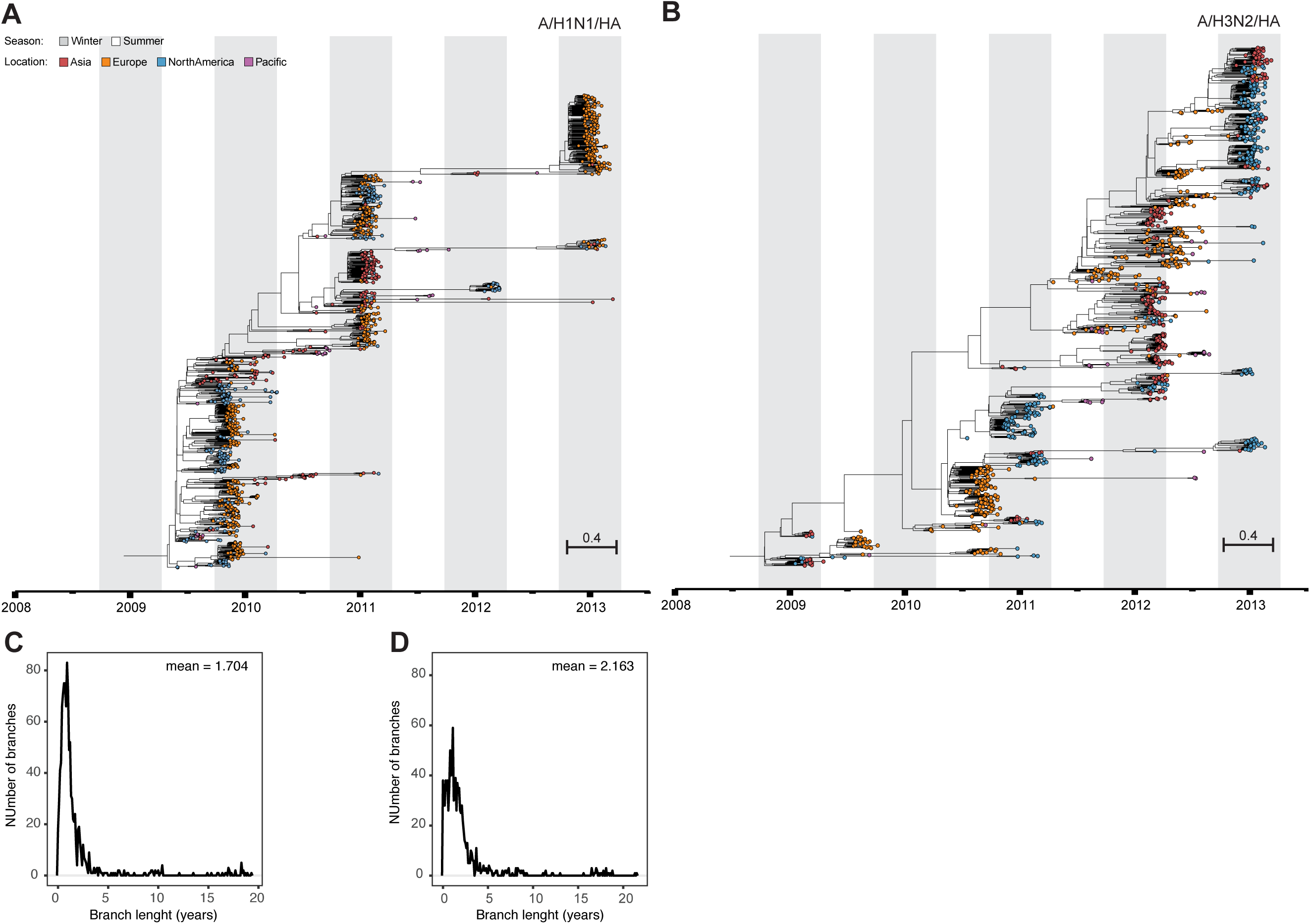
Spatio-temporal resolved phylogenies reveal intrinsic evolutionary influenza dynamics. Influenza hemagglutinin tree inferred by birth-death skyline (BDSKY) phylodynamic modelling using 1,078 (A/H1N1) (A) and 1,033 (A/H3N2) (B) complete gene sequences. Distribution of average trunk-to-tips branch lengths of A/H1N1 (C) and A/H3N2 (D) phylogenetic trees.

Besides these long-lived and predominantly strain-specific antibody-mediated immune responses, a shorter-lived, non-specific component has been proposed in particular to explain the limited virus genealogical diversity (single trunk) and lifespan (short-lived branches) of the vast majority of circulating dead-end virus lineages (11). Evidence for such component has first come from *in vitro* and animal studies showing that pre-infection with one subtype induces partial cross-protection from infection with another subtype (17). Observational studies addressing the potential role of cross-reactive immunity in global influenza seasonality has so far failed to show a clear pattern (18, 19). However, recent and marked observations to support a major role of cross-immunity were related to the fast disappearance of the A/H1N1 subtype from 1977, shortly after the introduction of the pandemic influenza A/H1N1 virus in 2009, while the A/H3N2 subtype managed to continue its circulation (13).

## Results

In search for the existence of such component we first followed a phylodynamic approach to jointly resolve spatio-temporal phylogenetic HA trees of A/H1N1 and A/H3N2 subtypes and to infer underlying host population dynamics (20, 21) (Fig. 1; Table S1 and S2). The dataset used here had been collected globally during the first 5 years after the onset of the 2009 influenza pandemic. It enrolled patients year-around (>1 year of age), the vast majority (>97%) with uncomplicated and PCR-confirmed influenza, who had been admitted - within 48 hours after symptom onset - to primary care centres and hospitals in Asia (Hong Kong; *n* = 6), Europe (*n* = 37), the US (*n* = 36) and the Pacific (Australia; *n* = 8) (Fig. S1). From these samples 2,111 influenza A viruses were isolated, which allowed us to obtain complete HA sequences from 1,078 A/H1N1 and 1,033 A/H3N2 viruses. The extent of sampling, directly after the pandemic outbreak, in combination with an unprecedented resolution regarding quality-controlled Sanger sequencing linked to patient data, resulted in a high-resolution dataset. This offered us a unique window of opportunity to study the dynamics of the estimated effective reproductive number (*R*) over time (*R*-skylines). *R* is a parameter of host immunity (22), and is computed here as the rate at which an infected individual gives rise to a new infection in a defined period of time.

We observed that *R*-skylines estimated from the A/H1N1 and A/H3N2 trees showed alternate phases of increasing and declining *R,* with *R* <1 and *R* >1 respectively (Fig.2). There was a significant negative correlation between phases (Pearson’s ρ = −0.511, *P* = 3.0e-07; D = 0.202, *P* = 0.052) with an average endogenous oscillation period estimated to be approximately 1.67 ± 0.01 years for A/H1N1 and 1.13 ± 0.02 years for A/H3N2 (6) (Fig. S2). Of note, these periods were similar to the average lifespan of the dead-end virus lineages on the HA trees (1.7 ± 0.4 for A/H1N1 and 2.2 ± 0.5 for A/H3N2) (Fig. 1).

**Fig. 2.**
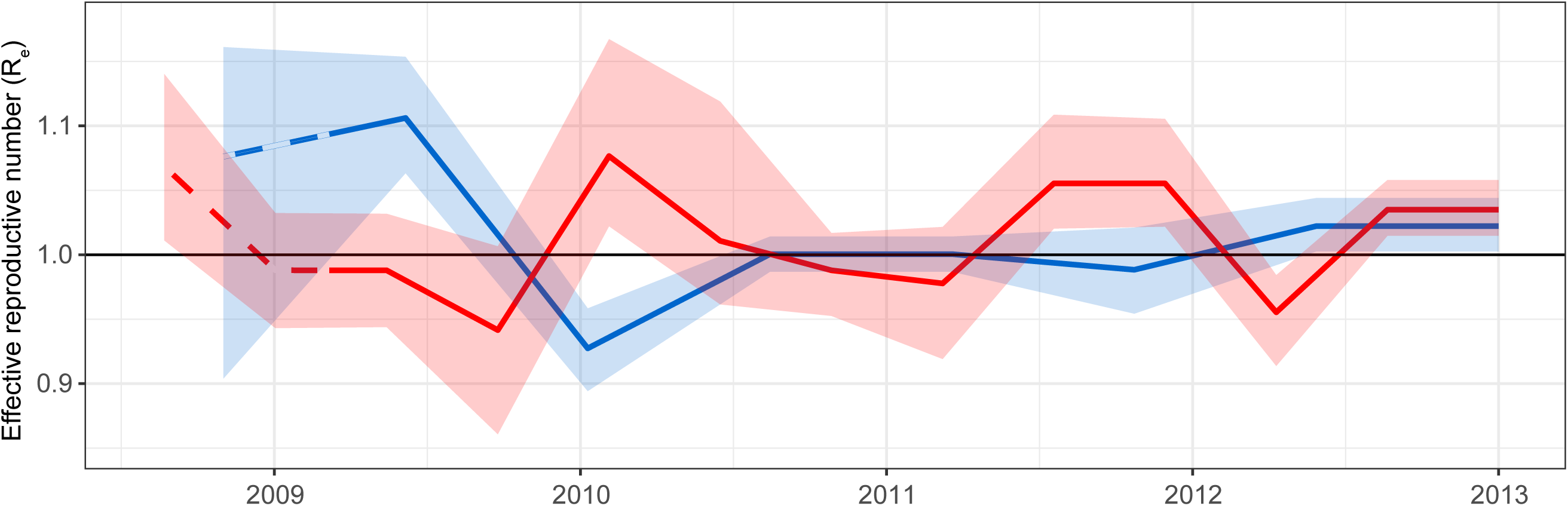
Oscillation of R-skylines estimated from influenza A virus phylogenies with opposed alternation of phases between subtypes. Time-series (2009-2013) for influenza A/H1N1 (blue, *n* = 1,078) and A/H3N2 (red, *n* = 1,033) viruses. Pre-pandemic period is indicated with dashed lines. Shaded regions represent 95% HPD interval.

Given the finite nature of susceptible hosts, virus persistence relies on the availability of new susceptible ones, which forces viruses to migrate between geographical locations (23). The interplay between antigenic drift and pre-existing immunity may then determine the outcome of the competition between these viruses at the onset of each influenza season (24). As the observed pattern of *R*-skylines indicates that a relatively short-lived cross-reactive immunity component exists, we wondered whether this competition also could determine the dynamics of global migration. To build on existing global migration data and to study its patterns for the A/H1N1 virus after 2009 we expanded our dataset with complete HA sequences deposited in the Influenza Resource Database (IRD) from viruses isolated prior to (2008-2009) and after (2013-2015) our study period (Dataset S1). We then inferred the number of geographical location changes at each internal node of these trees to identify all virus movements from one geographical location (source) to another location (sink). Again, and similar to the *R*-skylines, influenza A/H1N1 and A/H3N2 virus migration events alternated globally within our study period (Fig. 3). Global influenza A/H1N1 virus migration dominated in the first half of the study period, while A/H3N2 virus migration events were more prevalent between 2012 and 2013. This observation supports the evidence that inter-subtype competition presented here, contributes to influenza seasonality and may determine the virus that will dominate in a given influenza season.

**Fig. 3.**
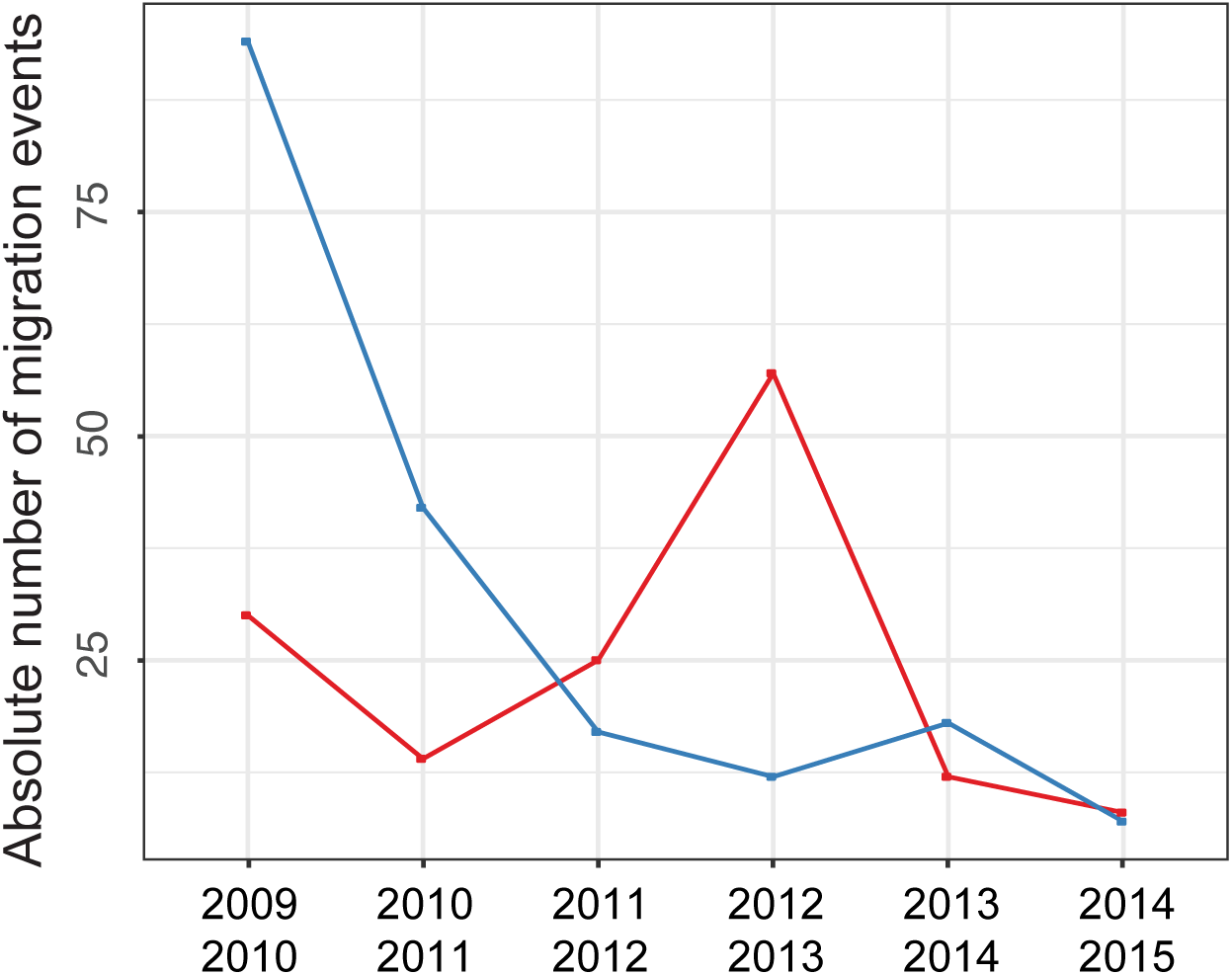
Opposed alternation of influenza A virus migration events. Total number of migration events between different geographical locations of influenza A/H1N1 (blue, *n* = 190) and A/H3N2 (red, *n* = 146) viruses were pooled by 1-year intervals from the start of the 2009 pandemic. Migration counts were performed by traversing the fully-spatiotemporal-resolved phylogenetic trees in post-order.

Finally, previous work on global circulation had shown that East and South-East Asia (E-SEA) played a pivotal role in global dissemination of A/H3N2 viruses. Here, A/H3N2 virus activity was found year-round (between 2000 and 2012), from where new antigenic drift variants fuelled in the temperate climate zone epidemics (9, 25). In contrast, E-SEA did not seem to have a major role in the dissemination of pre-pandemic A/H1N1 viruses (9). To study global virus migration after 2009 we constructed the networks of migration trajectories between the sampled geographical locations using a 1-year time window (Fig. 3 and 4; Fig. S3) and found that, in contrast to the pre-pandemic period (9), E-SEA was equally important for the dissemination of both influenza A viruses. Within our dataset we counted 56 A/H1N1 and 58 A/H3N2 dissemination events from E-SEA to the other sampled regions in the world (Fig. S4) In addition, global virus migration patterns showed a similar degree of global network complexity (Fig. S3, max. graph density/diameter of 1.08/12.09 for A/H1N1 and 1.16/10.64 for A/H3N2) and similar patterns of virus circulation across the sampled geographic regions (Fig. S3, max. number of islands and graph reciprocity of 2 and 0.75 for A/H1N1 and 2 and 0.86 for A/H3N2).

**Fig. 4.**
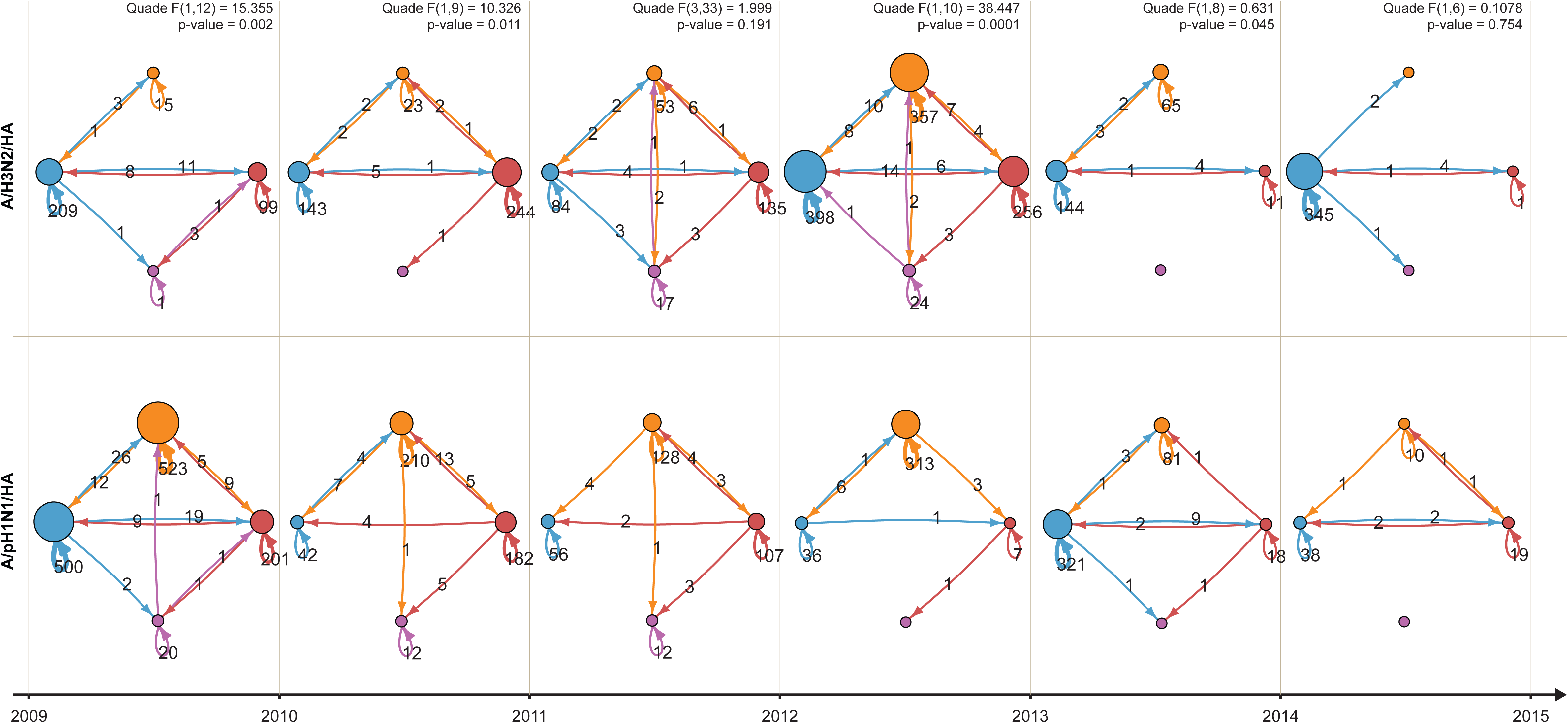
Network reconstruction of virus migration between the sampled geographic locations. Inferred migration networks between geographic locations. Centers located in Asia (red), Europe (orange), North America (blue) and Pacific (purple). Migration events during a 1-year time window were pooled. The diameter of the nodes is proportional to the number of sink migration events, while the arrow width is proportional to the number of source migration events. The Quade test and correspondent *post-hoc* procedures were applied to test significant differences between migration trends and preferred migration trajectories. Significance level was set to 5%.

## Discussion

Extrinsic factors probably play a role in forcing influenza epidemics into the winter seasons in the global temperate climate zones (3-5, 7). The oscillating and alternating pattern of the global skylines of *R* we present here indicate that cross-reactive host immunity is an important intrinsic driver of influenza seasonality (Fig. 2). Global influenza dissemination dynamics also reveals alternation of global virus migration events and complexity of migration trajectories between subtypes with two phases (Fig. 3 and 4). In the first phase (2009-2011) we observe that these parameters are high for the A/H1N1 and low for the A/H3N2 subtype (Fig. 3 and 4), but low for A/H3N2. This pattern is reversed during the second phase (2011-2013). These parameters are indicators of virus persistence and depend, therefore, on the availability of susceptible hosts within a defined geographical location (23). This implies that an intrinsic correlation exists between change of cross-reactive host immunity landscapes and global virus migration.

The observed short endogenous oscillation periods observed (1.7 years for A/H1N and 1.1 years for A/H3N2) suggest that this is most likely the result of short-lived inter-subtypic immune responses rather than antibody-mediated immunity from which major antigenic drift variants arise every few years (Figure 2 and S2) (9, 11, 16). Inter-subtypic immunity mediated by (CD4^+^ and CD8^+^) T-cells, and B-cells that generate or that trigger antibody-dependent cell-mediated cytotoxicity (ADCC) are proposed short-lived cross-reactive mechanisms (26-28). Taking into account the strong cross-reactive immune responses previously observed between the seasonal and pandemic A/H1N1 virus subtypes, cross-reactive immunity may well have forced the 2009 pandemic into the NH-summer and mitigated the disease burden associated with this virus (27, 28).

Further studies elucidating the contribution of host immunity to seasonality of influenza and other multi-strain viruses, such as the paramyxoviruses RSV and HMPV are warranted (29). This would further support the establishment and exploitation of global virus and serum banks (30), which will lead to a better understanding of the contribution of host immunity landscapes to the dynamic epidemiological circulation patterns of (multi-strain) pathogens.

## Materials and Methods

### Study conduct

IRIS (NCT00884117) is a prospective, multicentre, global observational study offering unprecedented resolution with regard to quality-controlled Sanger sequencing linked to patient data (31, 32). This report summarizes the results from 87 centers in, Australia (*n* = 8), China (Hong Kong, *n* = 6), Europe (*n* = 37; France, Germany, Norway) and the United States (*n* = 36) from December 2008 to March 2013, comprising five Northern and four Southern Hemisphere seasons, and including the 2009–pandemic. Centers were selected to achieve the widest geographic coverage possible within each country (Fig. S1). The study was performed in compliance with the principles of the Declaration of Helsinki and its amendments, and in accordance with Good Clinical Practice. Independent ethics committees and institutional review boards at each centre approved the study protocol and amendments.

### Patient selection

Adults and children aged ≥1 year were included year-round (*n* = 2980; excluding 21 patients (1%) with mixed influenza A and B virus infections) in the study if they were influenza-positive by rapid test (QuickVue Influenza A + B Test; Quidel Corp) at presentation and/or had predefined clinical signs and symptoms of influenza for ≤48 hours for hospitalized adults; ( ≤96 hours for hospitalized adults; no time limit for hospitalized children). The vast majority (> 97%) had uncomplicated influenza. All patients or legal guardians provided written informed consent at the time of enrolment.

### Assessments

Throat and posterior nasal swab specimens were obtained on day 1, 3, 6 and 10 and shipped on dry ice to a central laboratory for analysis (Erasmus MC, Rotterdam, the Netherlands). Influenza A subtypes types were identified using semi-quantitative real-time reverse transcription polymerase chain reaction (RT-PCR) (33). Day 1 samples with cycle threshold (Ct) values of < 32 were cultured on Madin-Darby canine kidney cells as described^4^. Virus-containing supernatants were cleared from cell debris by centrifugation (10 minutes at 1000 x g) and stored at −80°C until further processing. For this study, A/H1N1 (*n* = 1,078) and A/H3N2 (*n* = 1,033) virus isolates were included, which were obtained at patient admission (day 1).

### Datasets and nucleotide sequence accession numbers

Sanger sequencing of hemagglutinin (HA) genes was done for all isolated viruses. Complete HA sequences were obtained for influenza A/H3N2 (*n* = 1,033; gi:XX12345-XX12345) and A/H1N1 (*n* =1,078; gi:XX12345-XX12345) subtypes. To build on existing data on global influenza migration we expanded the IRIS dataset with all available complete HA and NA sequences from the NIAID Influenza Research Database (IRD) collected between 2008-2009 and 2013-2015 in countries included in the IRIS study (34). Numbers of additional HA sequences were 443 for A/H1N1 and 462 for A/H3N2 respectively. The complete list of IRD sequences is provided (Table S1).

### Data pre-processing and alignments

Each expanded dataset was aligned using ProGraphMSA using default parameters (35). Sequences were renamed to include sampled geographical locations, sampling dates (continuous values) and corresponding influenza season (when available).

### Phylodynamics inference

The BDSKY phylodynamics model implemented in BEAST v.2.3.1 was applied to the IRIS and expanded datasets to infer spatio-temporal resolved phylogenies and epidemiological parameters (20, 23, 36). Phylogenetic trees were estimated under the general-time-reversible model (GTR+Γ_4_) with Γ-distribution to model among-site rate variation (37)(Fig. 1). A molecular clock rate prior was set to follow an uncorrelated log-normal distribution (38). Internal node calibration was performed using tip sampling dates (39, 40). The BDSKY-model was set with the following parameter: The sampling rate prior for the influenza infected population/the real sampled population followed a *Beta (1, 999)* distribution; the prior probability of sampling an individual upon becoming non-infectious followed a *LogNorm (4.5, 1.0)* distribution. In addition, tree dating was performed using tip dates while an uncorrelated log-normal clock rate prior was applied to handle uncertainties in the sample collection dates. Finally, the analysis was run long enough to obtain a sufficient effective sample size ESS > 200 for all parameters. The converged parameters of the BDSKY-model are listed in supporting information table S1 and S2. To assess global model robustness, we performed two independent runs of each analysis (for a total of 20 runs). MCMC parameter convergences were diagnosed with Tracer 1.6. Thinning of BEAST2 output files (tree files and parameter files) was done using in-house bash scripts. After accurate MCMC trace monitoring, the first 10% of MCMC steps were discarded as burn-in resulting in around 6000 trees per each dataset. TreeAnnotator v2.3.1 was used to produce Maximum Clade Credibility (MCC) trees (20).

### Statistical analyses Effective reproductive number

We estimated the effective reproductive number *R* using phylodynamics modelling as described above. The estimates of *R* allowed us to study the dynamics of virus spread within the population (41). Values *R* < 1 indicate a decline of infections, while *R* > 1 indicates that the infection has increased its spreading in a more susceptible population. The skyline of *R* is used here to picture the underlying dynamics ‘shaping’ a phylogenetic tree (20, 21, 23). The univariate distributions of *R* values, estimated with independent sampling frequency from each dataset, were grouped and smoothed via interpolation to compensate for intermediate missing values. The dataset was set to start from a sampled common date (2009.24508). First, Wald-Wolfowiz, and Bartel Rank non-randomness tests were applied on each *R* median time-series as well as its permuted version (42). The same test was then applied on the pairwise intersection of *R* median time-series, and the statistical support was evaluated by re-computing the test on permuted *R* median time-series. Secondly, the pairwise maximum difference between *R* median time-series was computed applying the Kolmogorov-Smirnov test (KS-test). The two-sample Kolmogorov-Smirnov test was used to compare the cumulative distributions of two data sets (43). The KS-test reports the maximum difference between two cumulative distributions (D) and it returns a *P* computing the KS statistics from all the possible permutation of the original data. The significance level was set at 0.001, so that two distinct *R* median time-series were considered to be drawn from different distributions when D ≥ 0.45. Next, the pairwise-correlation between *R* median estimates was evaluated by the Pearson's product moment correlation coefficient (*ρ*). Pearson’s product moment correlation coefficient (*ρ*) was tested using the Fisher’s Z transform with 95% confidence interval and significance level set at 0.005 (44). Exploratory analyses on the *R* median time-series were applied to qualitatively identify oscillation periods and amplitude. The oscillation period of each *R* median time-series was then computed from the highest frequency value shown by the smoothed periodogram using the IRIS dataset. Statistical uncertainty on the inferred period was assessed from cumulative periodograms computed on 100 permutations of the original *R* median time-series. Finally, the overlap of HPD intervals of the pairwise *R* was computed for each R median time-series. The obtained value was then compared with the overlap of the HPD interval of *R* obtained with 100 permutations of the true HPD intervals.

### Migration routes and evolutionary rates

Datasets were partitioned according to the geographical sampling locations pooled by continent (North America, Europe, Asia, Pacific area). Migration rates were estimated using a discrete phylogeographic trait model with the Γ-distribution as substitution rate prior between geographical demes (45, 46). The influenza virus dissemination process was fitted to a discrete trait model using the Bayesian Stochastic Search Variable Selection (BSSVS) method, by inferring the most parsimonious description of the phylogeographic diffusion process (47-50). Counts of migration events were quantified by traversing the fully spatio-temporal resolved phylogenetic trees in post-order and by counting the number of most probable Markov chain jumps along the branches of the posterior set of trees (PSTs) (51, 52).

### Branch geographical persistence

Geographical persistence was quantified by summing the phylogenetic branch lengths (measured in expected substitutions per site) grouped by their inferred geographical location on the phylogenetic tree trunk (inferred traversing the phylogenetic tree from ‘leaf-to-root’ and summing the number of branch traversals. The tree trunk was defined as the path on the phylogenetic tree that has been traversed more than 10 times. Number of seeding events was defined as the number of switches on the phylogenetic tree trunk per season.

### Migration graphs

Trajectory networks were reconstructed per each strain variant, pooling migration events occurred within a one-year time window (Fig. 4). Trajectory complexity was computed estimating graph density, number of islands (nodes), diameter, and reciprocity (53, 54). In addition, geographical location connections were estimated by computing the graph centrality measures (specifically: degree centrality and betweenness centrality) (Fig. S3 and S4) (55-57). The Quade and correspondent *post-hoc* procedures were applied to test whether migration trends were significantly different between strains and whether preferred migration trajectories were selected (58). The significance level was set at 5%.

### Data availability

Full-length HA sequences obtained as part of the IRIS study have been deposited in Genbank with the primary accession codes mentioned above. All other HA sequences downloaded for this study are listed in the Supportive information.

### Code availability

All source codes and BEAST .xml files are available on GitHub (http://www.github.com/gattil/IRIS-Influenza-Dynamics).

## Acknowledgments

We thank Christian von Mering, Urs Greber and Alexander Roth (University of Zurich) for constructive inputs during discussions. Tanja Stadler, Denise Kühner, Veronika Boskova (ETH Zürich) and Alexei Drummond (University of Auckland) helped at an early stage of the analysis with the BEAST 2 software. We thank Anne Van der Linden, Martine Bakx, Danielle de Wijze, Jolanda Maaskant, Jeer Anber, Moniek van der Haven of the IRIS-team and Be Niemeijer from ErasmusMC for their assistance in the lab and Martin Ebeling from Roche for help on sequence analysis. We thank Xiao Tong, Isabel Najera, Klaus Klumpp, James Smith and Laurent Essioux for insightful discussions and project support. All collaborating physicians at the participating centers are thanked for their support at the sites and for sending in the samples.We thank MICRON research for the global logistics. Special thanks to all the patients who were willing to participate in the IRIS trial. LG is funded by SNSF Projektförderung no. 157064 and he was granted with UZH-ETH MLS PhD Program Travel Grant. This research was supported by the Volkswagen Foundation.

## Author contributions

AO and MS participated in the design of the IRIS study. EV and MS collected the data. LG and MA designed the analysis pipeline. LG implemented the software pipeline and analysed the data. EV, LG, JDZ, MA and AO interpreted the results. EV, LG, MA, JDZ and AO wrote the paper.

---

**Fig.S1.**
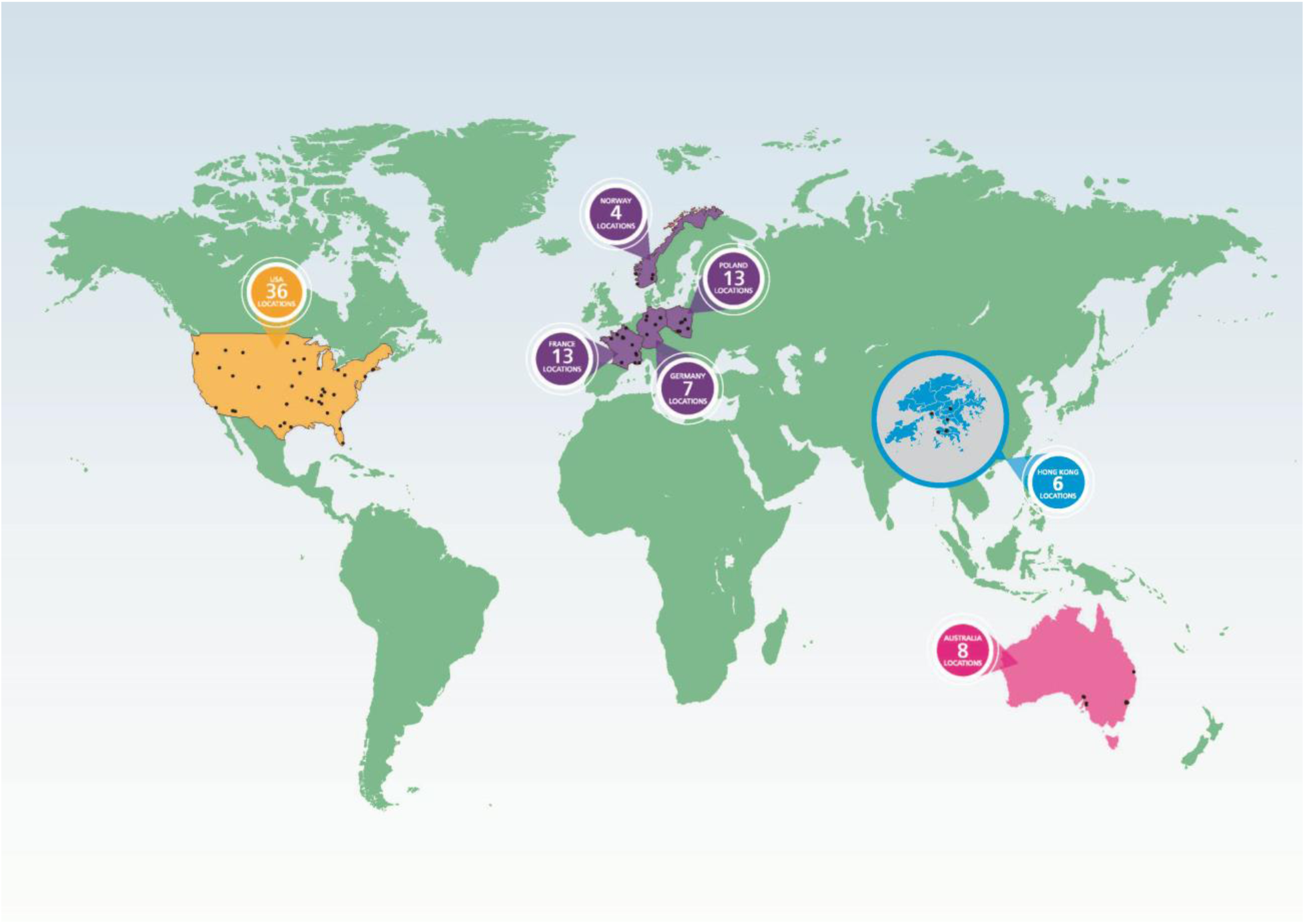
Distribution of IRIS centres over the world. IRIS (NCT00884117) is a prospective, multicentre, observational study. This report summarizes the results from 87 centres in, Australia, China (Hong Kong), France, Germany, Poland, Norway, and the United States from December 2008 to March 2013, comprising five Northern and four Southern Hemisphere seasons, and including the 2009–2010 pandemic.

**Fig.S2.**
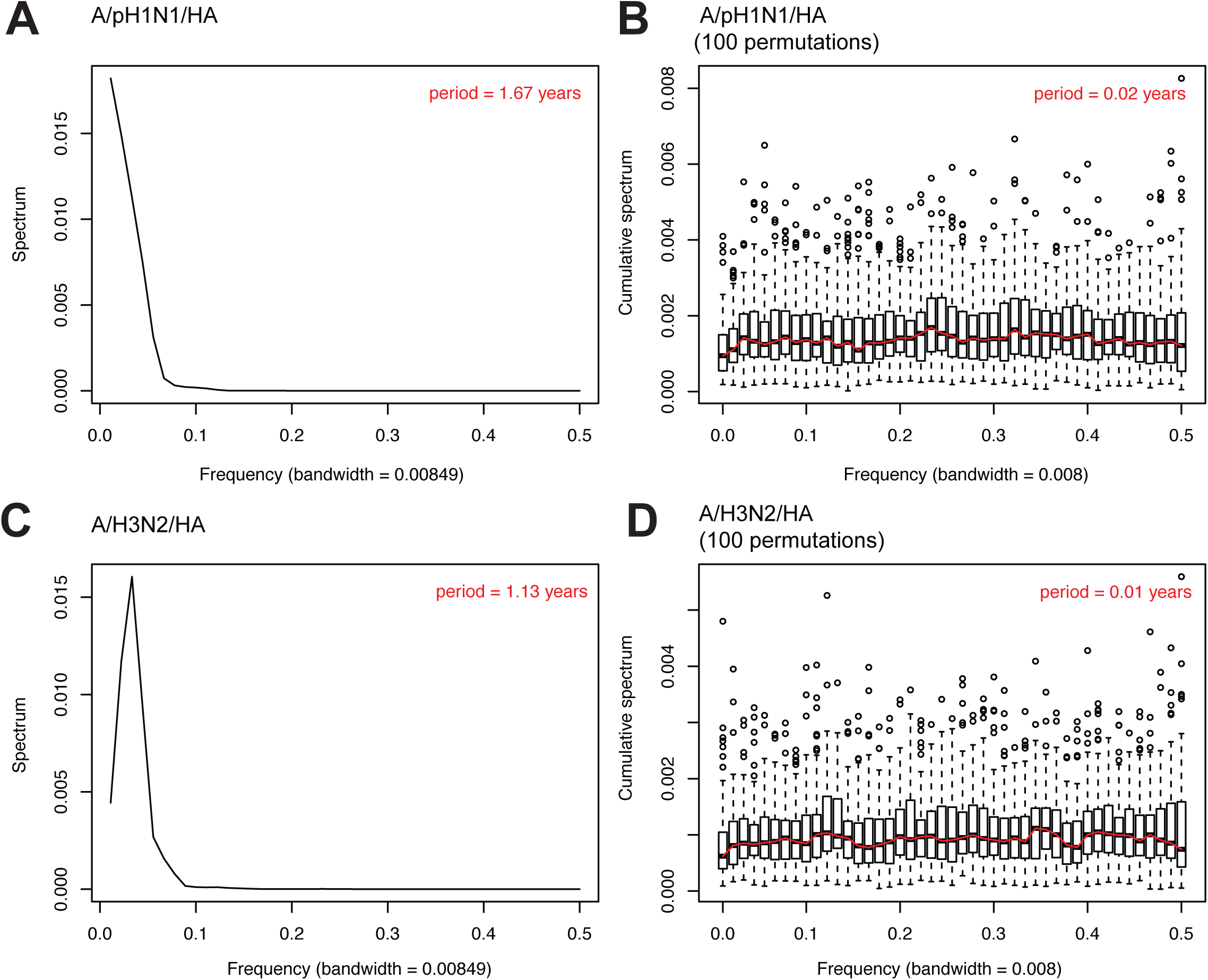
Periodograms computed for the *R*-skylines to estimate oscillation periods and inferred uncertainties. Estimated periods for the A/H1N1 (A) and A/H3N2 (C) R-skyline plots. Uncertainty quantification on estimated periods for A/H1N1 (C) and A/H3N2 (D) virus was done using cumulative periodograms computed on 100 permutations of the original *R* median time-series.

**Fig.S3.**
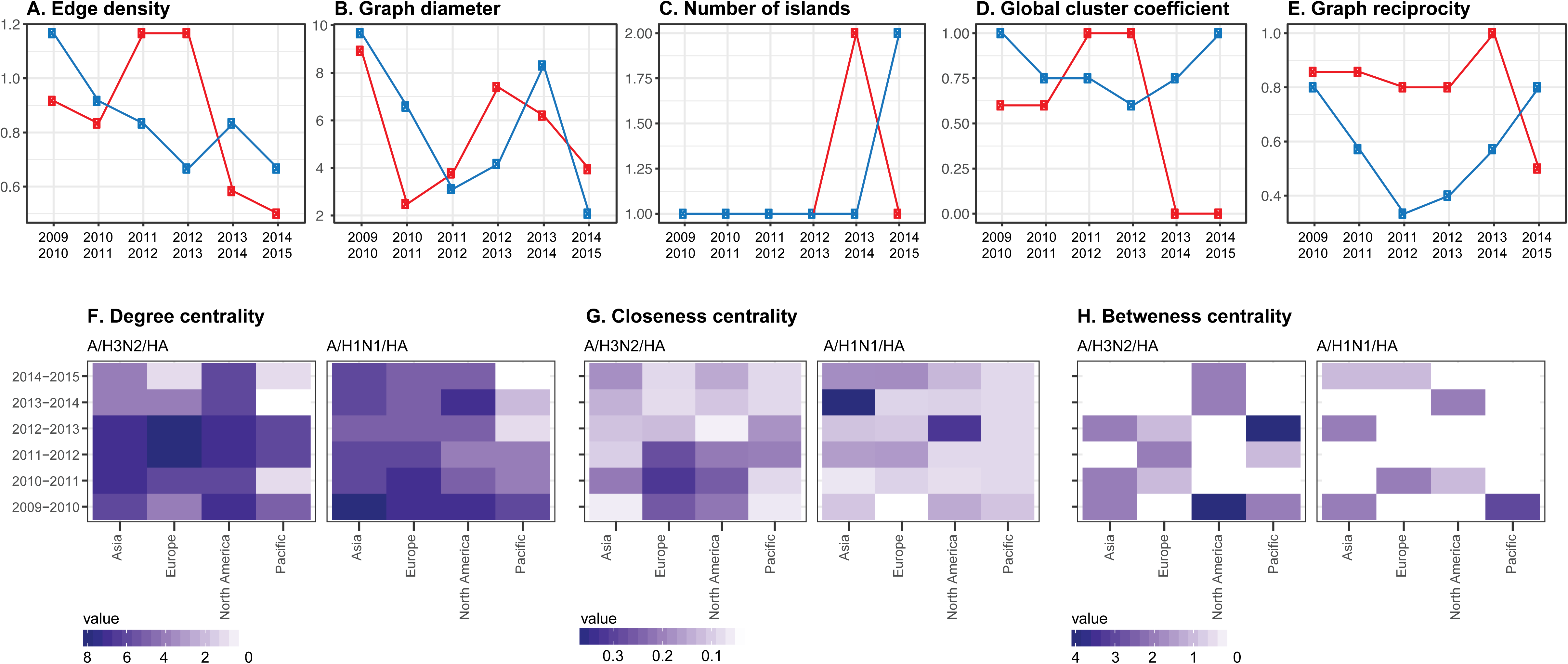
Migration network measures and clustering analyses of reconstructed migration networks Complexity of network connections was quantified for A/H3N2/HA (red) and A/pH1N1/HA (blue) using the following measures: (A) Network density. (B) Graph diameter.(C) Number of islands. (D) Global cluster coefficient.(E) Reciprocity of the graph. Comparison of the connectivity between different geographical locations was done within a 1-year-time-window frame using the following measures: (F) Degree of centrality. (G) Closeness centrality. (H) Betweenness centrality.

**Fig. S4.**
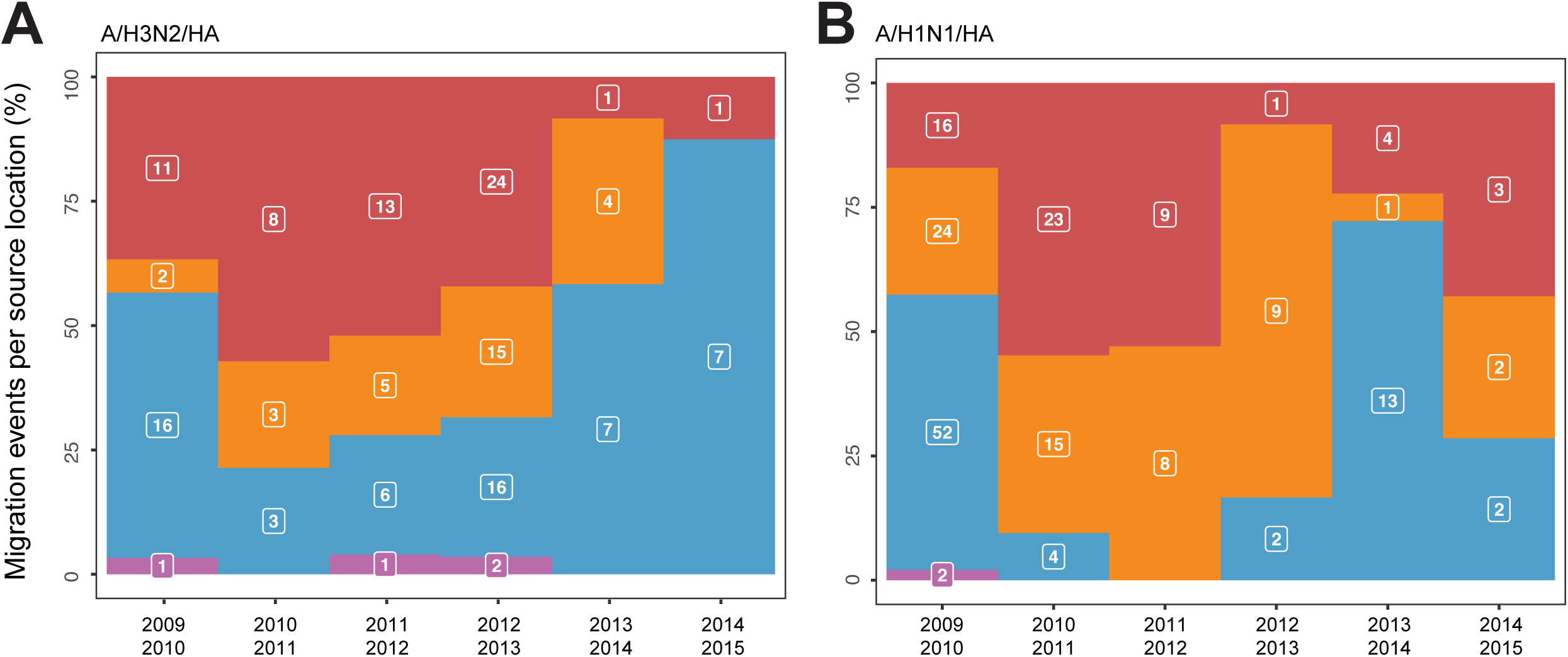
Distribution of migration events between geographical locations. Influenza A/H3N2 (A) and A/H1N1 (B) virus migration events were pooled by 1-year intervals. Counts of geographic location switches on the tree were identified and classified using fully-spatiotemporal-resolved HA phylogenies by a discrete source/sink model between centers located in Asia (red), Europe (orange), North America (blue) and Pacific (purple) and are presented in white boxes.

